# IRESbase: a Comprehensive Database of Experimentally Validated Internal Ribosome Entry Sites

**DOI:** 10.1101/2020.01.15.894592

**Authors:** Jian Zhao, Yan Li, Cong Wang, Haotian Zhang, Hao Zhang, Bin Jiang, Xuejiang Guo, Xiaofeng Song

**Author notes:** Equal contribution (Jian Zhao, Yan Li, Cong Wang, Haotian Zhang). Corresponding author(s). (Guo X), (Song XF).

## Abstract

Internal Ribosome Entry Sites (IRESs) are functional RNA elements that can directly recruit ribosomes to an internal position of the mRNA in a cap-independent manner to initiate translation. Recently, IRES elements have attracted much attention for their critical roles in various processes including translation initiation of a new type of RNA, circular RNA, with no 5′ cap to support classical cap-dependent translation. Thus, an integrative data resource of IRES elements with experimental evidences will be useful for further studies. In this study, we present a comprehensive database of IRESs (IRESbase) by curating the experimentally validated functional minimal IRES elements from literature and annotating their host linear and circular RNA information. The current version of IRESbase contains 1328 IRESs, including 774 eukaryotic IRESs and 554 viral IRESs from 11 eukaryotic organisms and 198 viruses. As our database collected only IRES of minimal length with functional evidences, the median length of IRESs in IRESbase is 174 nucleotides. By mapping IRESs to human circRNAs and lncRNAs, 2191 circRNAs and 168 lncRNAs were found to contain at least one entire or partial IRES sequences. The IRESbase is available at http://reprod.njmu.edu.cn/cgi-bin/iresbase/index.php.

## Introduction

Translation initiation is a crucial and highly regulated step during protein synthesis in eukaryotes. Generally, eukaryotic mRNAs recruit ribosomes to initiate translation by a canonical cap-dependent scanning mechanism, which requires a methylated cap structure at the 5′ end of mRNAs and is mediated by the eukaryotic translation initiation factors (eIFs). Instead of the cap-dependent manner, a subset of eukaryotic mRNAs can initiate an internal translation through specialized sequences, called internal ribosome entry sites (IRESs), which can directly recruit 40S ribosomal subunits to initiate translation.

The IRES elements were first discovered about 30 years ago in picornaviruses [1], following the discovery of novel IRESs in many other pathogenic viruses (*e.g.*, EMCV, FMDV, and HIV) [2–4]. The IRES elements ensure the efficient viral translation, when canonical eIFs necessary for mRNA recruitment are inhibited in infected cells. Besides the viral mRNAs, IRESs were also found in a subset of eukaryotic mRNAs (*e.g., XIAP, p53, DAP5*, and *Apaf-1*) [5–8]. The IRESs in eukaryotic mRNAs allow their translation to continue under many physiologically stressful conditions when the cap-dependent translation is suppressed. In addition, for those eukaryotic IRES-containing mRNAs with long and highly structured 5′-UTRs (incompatible with the cap-dependent scanning mechanism), the IRES elements ensure their efficient translation under normal physiological conditions even when cap-dependent translation is fully active [9]. The increasing number of discovered eukaryotic IRESs suggests that IRES-driven translation initiation accounts for a significant proportion of eukaryotic mRNAs. Although the studies of IRESs have been limited, eukaryotic mRNAs bearing IRESs seem to play important roles in diverse processes, including stress responses, cell survival, apoptosis, mitosis, tumorigenesis and progression [10–14].

The two IRES databases, IRESdb and IRESite were built in 2002 and 2005, respectively [15,16]. In addition, the Rfam database collected IRESs as a type of cis-regulatory RNA element [17]. However, they only include IRESs in mRNAs. Recently, the IRES elements were also found in the circular RNAs (circRNAs) and long noncoding RNAs (lncRNAs) [18]. The circRNAs, *circ-FBXW7, circ-ZNF609, circ-SHPRH, circPINTexon2*, and *circβ-catenin*, were found to be translated into polypeptides mediated by IRES elements [19–23]. The translation of two small open reading frames (120 nucleotides and 141 nucleotides long) within the lncRNA *meloe* was achieved by an IRES-dependent mechanism [24]. Identification of the translation of circRNAs and lncRNAs driven by IRES elements has attracted more attention recently.

The IRESdb database only has 30 viral IRESs and 50 eukaryotic IRESs, and the IRESite database contains 125 IRESs from 43 viruses and 70 eukaryotic mRNAs. The Rfam database built 32 RNA families with about 11 viral IRESs and 21 eukaryotic IRESs. In this study, by manual curation, we developed a novel non-redundant public database and named it IRESbase. This database contains updated experimentally validated IRESs, including 554 viral IRESs, 691 human IRESs, and 83 IRESs from other eukaryotic species.

In order to facilitate development of new IRES-dependent functional regulation studies, we developed a new resource database named IRESbase which is rich with information about human IRESs. This database includes information regarding the genomic positions, sequence conservation, SNPs, nucleotide modifications, targeting microRNAs, host gene, host transcripts (mRNA, circRNA, and lncRNA), GO terms (biological process, molecular function, and cellular component), KEGG pathway annotations, and several critical experimental assay information. In the database, 2191 circRNAs and 168 lncRNAs were found to contain at least one entire or partial IRES element. Among these transcripts, 1012 circRNAs were found to contain an open reading frame (ORF) longer than 300 nucleotides, and all the 166 lncRNAs were found to contain at least two small ORFs (ranging from 60 to 300 nucleotides in length). Our IRESbase database allows users to search, browse, and download as well as to submit experimentally validated IRESs.

## Results

### Database content

IRESbase is intended to be a comprehensive database that includes both eukaryotic and viral IRESs with experimental evidences and covers not only the IRES sequences and structural information but also the IRES host transcript and gene information. The database currently contains a total of 1328 IRES entries, including 774 eukaryotic IRES entries and 554 viral IRES entries. Each eukaryotic IRES entry contains three basic types of records – IRES, gene, and transcript records if available, each viral IRES entry contains only the IRES and transcript records. In total, 1328 IRES records, 725 eukaryotic gene records, 3633 eukaryotic transcript (1274 mRNAs, 168 lncRNAs, and 2191 circRNAs) records, and 588 viral transcript records are included in our established IRESbase. The data sources and the structure of IRESbase are shown in **Figure 1**.

**Figure 1.**
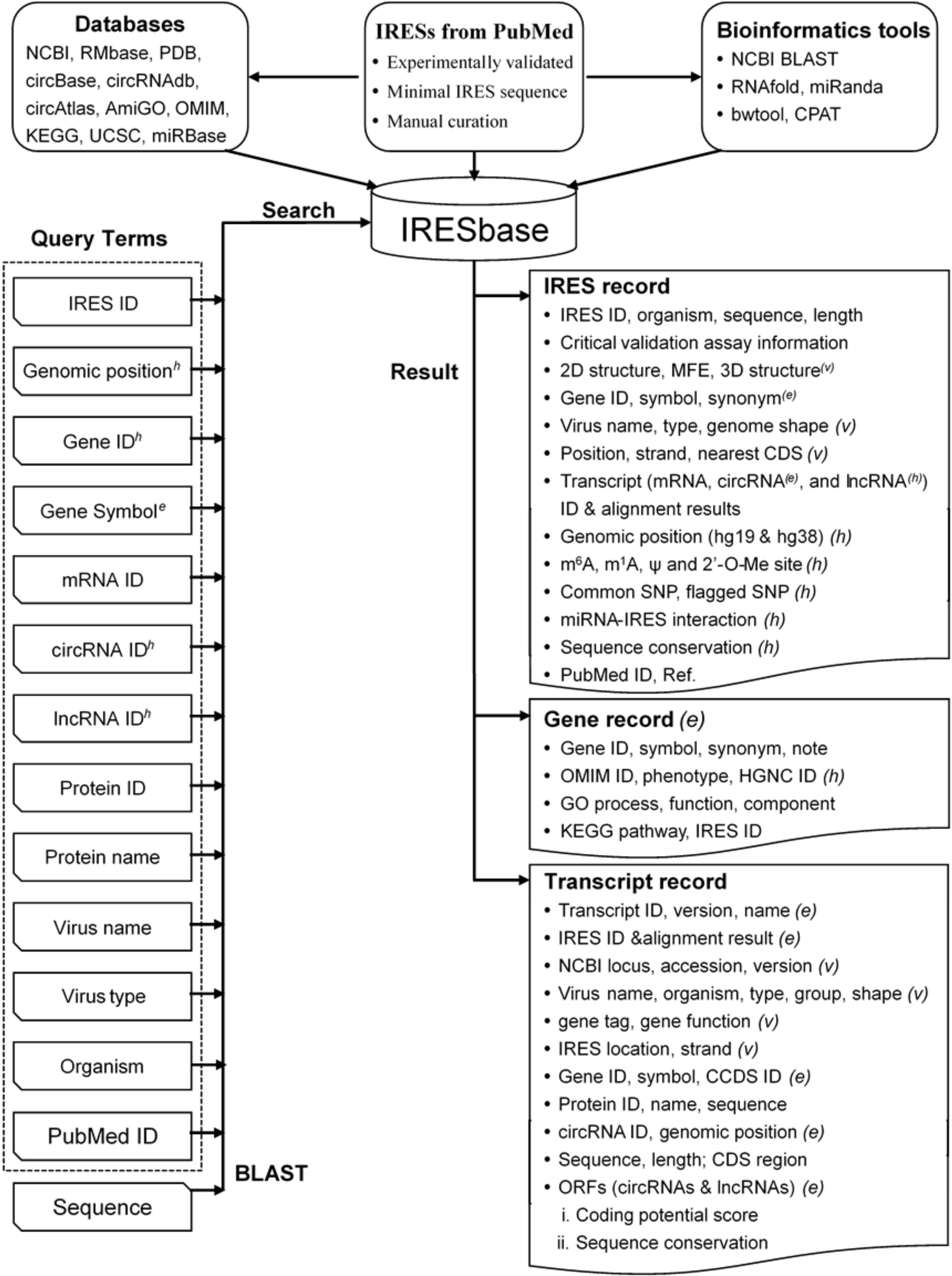
Data sources and the IRESbase structure. For record terms, the ‘*(e)*’, ‘*(v)*’, and ‘*(h)*’ tags indicate eukaryotic data, viral data, and human data, respectively. If the tag is located at the end it indicates the tag is shared by all the terms in the line, while the superscript tag indicates that the tag only belongs to the tagged term itself. Terms without tags are commonly used for different species. For query terms, superscript ‘*e*’ is exclusively for the eukaryotic data, superscript ‘*h*’ is exclusively for the human data, while the remaining terms without superscripts are commonly used in different species.

As the available data are different in different species, human IRES records have the largest number of attributes (32 in total, if available); the IRES records for other eukaryotic species and viruses have 20 and 24 attributes, respectively (Figure 1). The 32 attributes of the human IRES record are IRESbase ID, organism, sequence, sequence length, seven critical validation assay information (i.e., bicistronic backbone, positive and negative controls, second cistron expression, tested cell types, methods used to analyse RNA structure, and *in vitro* translation experiments), predicted two-dimensional (2D) structure, minimal free energy (MFE), verified 2D structure, PUBMED ID, reference, host gene ID, host gene symbol, host gene synonym, host mRNA ID and its IRES alignment result, genomic position (location and strand) in hg19, genomic position in hg38, host lncRNA ID and its IRES alignment result, host circRNA and its alignment result, sequence conservation, common single nucleotide polymorphism (SNP), flagged SNP, N6-methyladenosine (m^6^A) site, N1-methyladenosine (m^1^A) site, pseudouridine modification (ψ) site, 2′-O-methylation (2′-O-Me) site, and predicted microRNA-IRES interaction. The IRES record for other eukaryotic species only has the first 20 attributes, and the first 18 attributes are included in the viral IRES record. The viral IRES record contains six other attributes including viral genome ID (*e.g., NC_006273*), genome shape, virus type, genomic position (location and strand), coding sequence (CDS) of regulation, and three-dimensional (3D) structure.

As shown in Figure 1, the eukaryotic gene records commonly have nine attributes, which are gene ID, gene symbol, gene synonym, gene note, GO process, GO function, GO component, KEGG pathway, and IRESbase ID. For human, besides the above nine attributes, three other attributes (OMIM ID, OMIM phenotype, and HGNC ID) are included. The eukaryotic transcript records for IRES host mRNA, lncRNA, and circRNA contains 12, 13, and 11 attributes, respectively. The 12 attributes of eukaryotic mRNA record are transcript ID, name, version, sequence, sequence length, IRESbase ID & IRES alignment result, gene symbol, CDS region, protein ID, protein name, CCDS ID, and sequence. For lncRNA record, besides the first seven attributes in the mRNA record, the other six attributes are gene ID, ORF, ORF’s coding potential score, ORF’s sequence conservation, predicted protein sequence and length. For circRNA record, the 11 attributes are circRNA ID (circBase ID, circRNADb ID, and circAtlas ID) [25–27], genomic location, strand, gene symbol, best transcript, IRESbase ID and the IRES alignment result, ORF, coding potential score, predicted protein sequence, protein length, and reference. The viral transcript record has 21 attributes, including NCBI locus, accession, version, virus name, organism, virus type, virus group, transcript sequence, sequence length, transcript shape, gene name, gene ID, gene tag, gene function, CDS region, protein name, protein ID, protein sequence, IRESbase ID, IRES location, and strand.

### Database interface

#### Simple, advanced, and BLAST search

IRESbase was developed in a user-friendly mode, providing a search engine to find the IRESs of interest. The homepage (Figure S1) offers a search box for querying the IRESbase by IRES ID, gene ID, transcript (*e.g.*, mRNA, lncRNA, and circRNA) ID, protein ID, PubMed ID, chromosomal location, organism name, gene name, protein name or a substring of any of the above keywords. If a keyword (*e.g., FUBP1* or HCMV) is added into the search bar, the results will be shown in a tabular format. By clicking on the IRES ID (*e.g., hsa_ires_00089.1* or *vir_ires_00484.1*), the detailed information will be shown in three parts: the IRES information, its host gene information, and its host transcript information (**Figures 2 and 3**). Furthermore, the terms in the page are cross-linked to several external databases including NCBI, UCSC, AmiGO, KEGG, HGNC, RMbase, circBase, and circRNADb [25,26,28–32].

**Figure 2.**
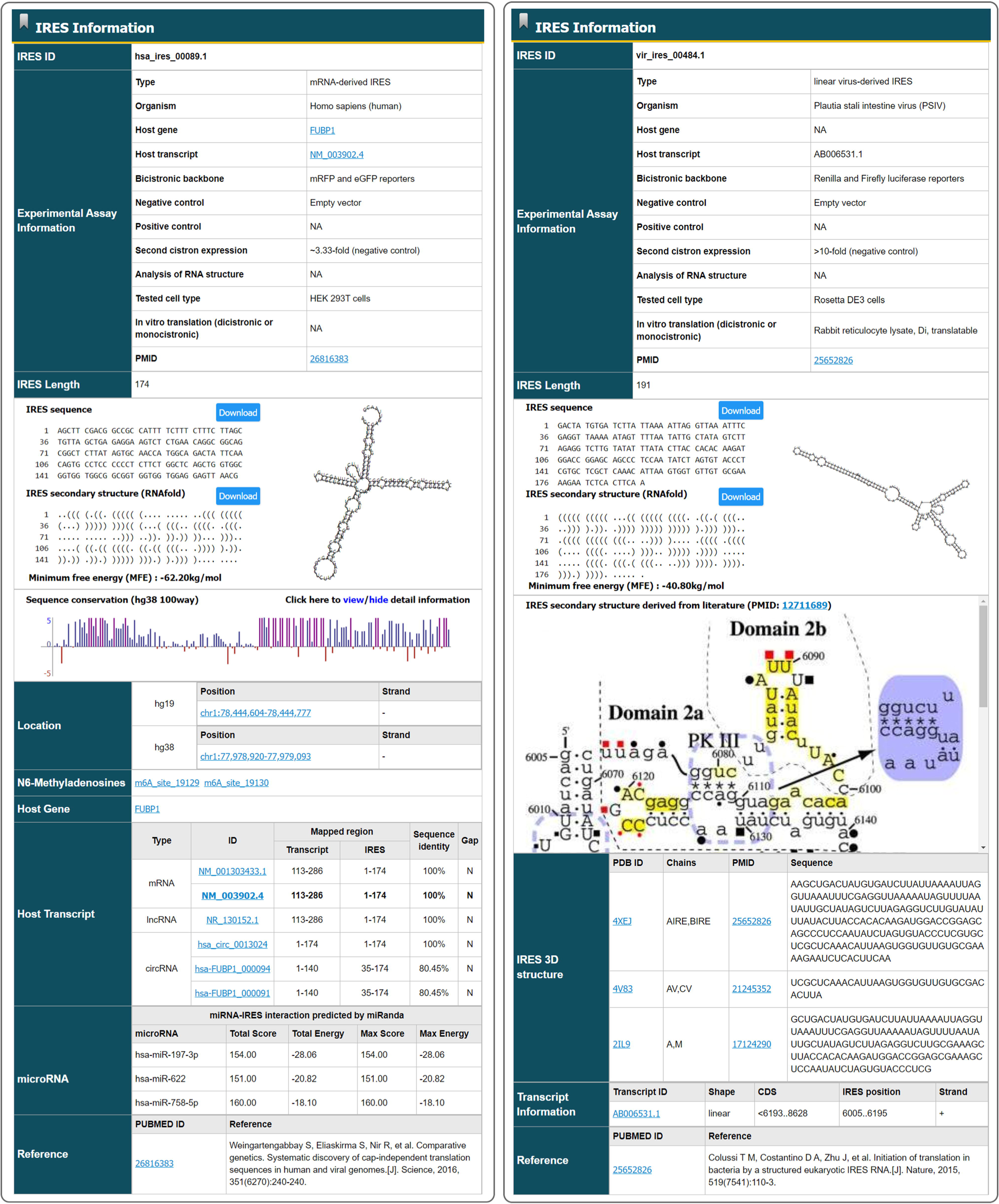
The IRES entry page (Left: *hsa_ires_00089.1*; Right: *vir_ires_484.1*) The terms shown in the ‘miRNA’ field of ‘*hsa_ires_00089.1*’ (left) were truncated for easy presentation.

**Figure 3.**
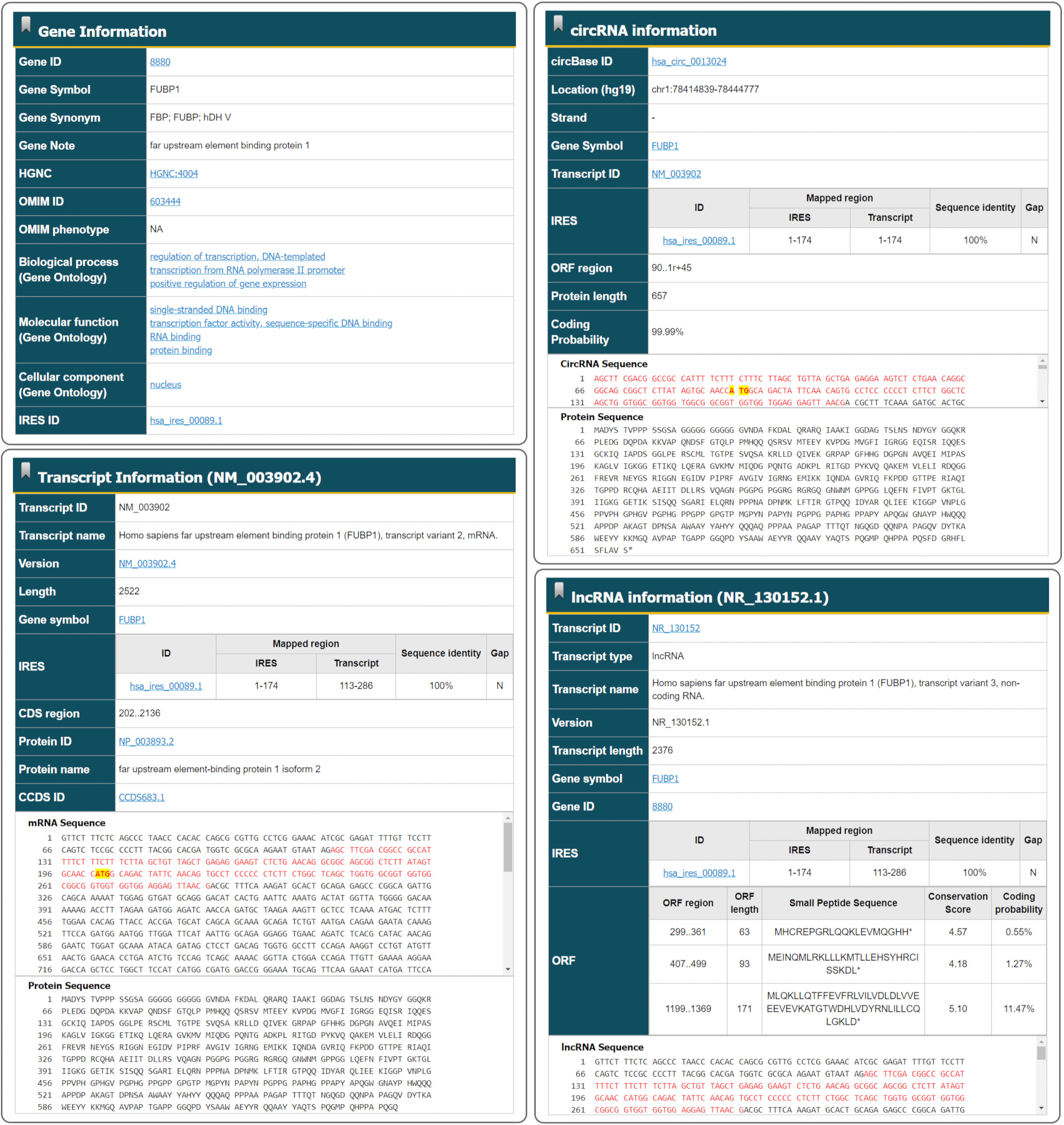
The gene and transcript (mRNA, lncRNA, and circRNA) records in the page of a human IRES entry (*hsa_ires_00089.1*) The IRES region is marked in red and the boundary codons of the CDS region are highlighted with yellow.

In addition, IRESbase provides three additional advanced search options for users, including (1) ‘BLAST search’, (2) ‘Advanced search’, and (3) ‘Chromosome location’.

1. BLAST search: this option was designed to find if there was a similar IRES in the RNA sequence of interest. It provides four IRES sequence datasets (*e.g., Homo sapiens*, other eukaryotic organisms, virus, and all organisms) for users. Users can input more than one query sequence in FASTA format. The search results would be summarized in a table containing job title, IRES ID, score, expect, identity, gaps, and strand (**Figure 4**). By clicking the ‘Job title,’ the alignment detail will be shown at the nucleotide level.
2. Advanced search: in this option, users can use one or more query fields together to retrieve the IRESs of interest. Different query fields are provided for users to search IRESs of different organisms, including virus, *Homo sapiens*, and other eukaryotic organisms (Figure 1).
3. Chromosome location: this option was designed for users to find if there was a verified IRES located in the circRNAs of interest through the input of a specific chromosomal region in *Homo sapiens* (hg19 or hg38).

#### Data browse, download, and submission

Instead of searching for a specific IRES, all entries of IRESbase were grouped by organism, GO & KEGG, chromosome, and transcript type. Users can browse the IRES group of interest by clicking the count on the left of the corresponding page. Unlike other existing IRES databases, IRESbase supports users to download all the IRES records together in a simple tab-delimited format or to download each IRES record separately in a standard PageTab (Page layout Tabulation) format, which only contains several basic IRES information and the validation assay information. In addition, users can also select those IRES attributes of interest to download in an Excel-compatible format.

**Figure 4.**
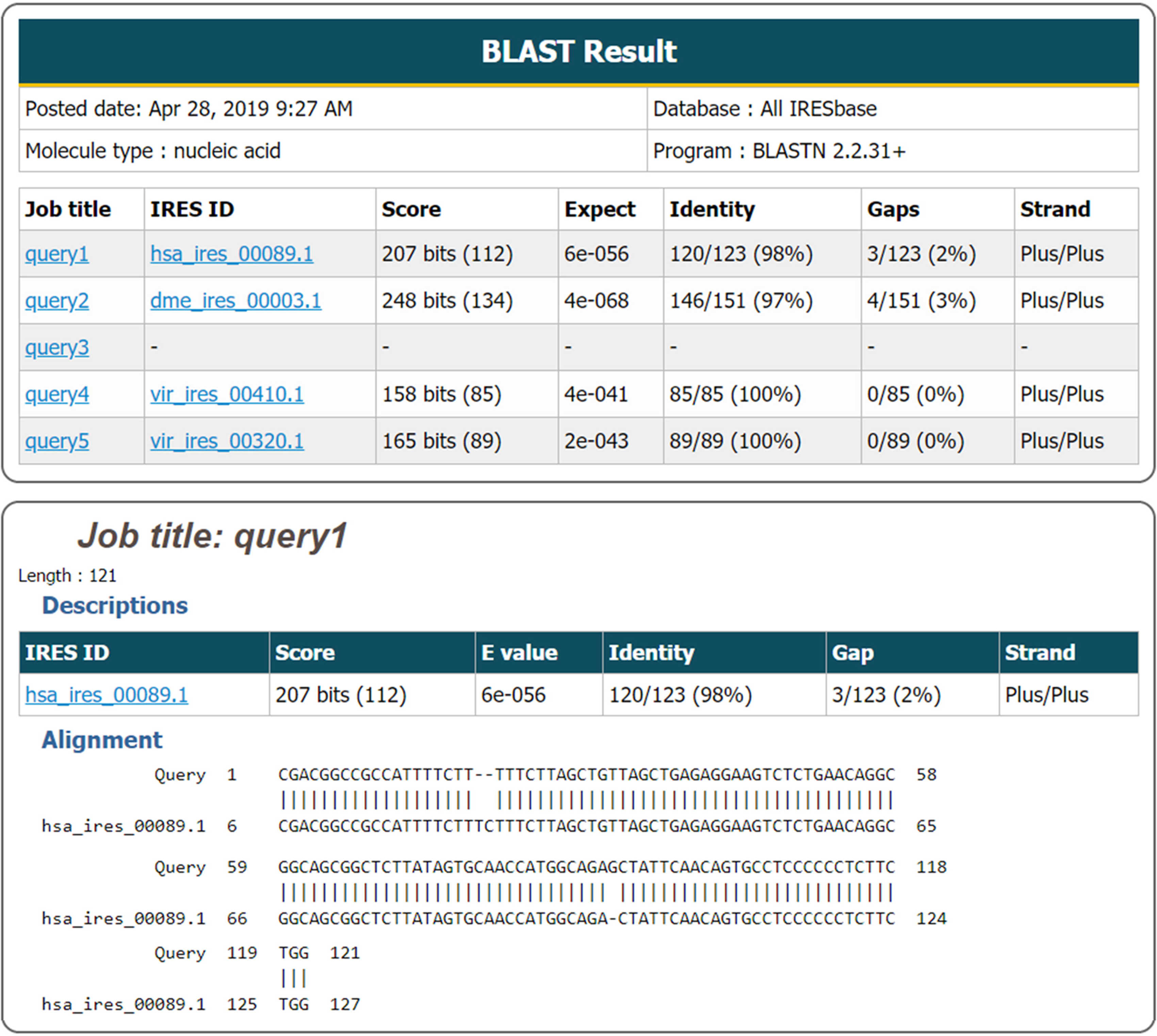
The result page of BLAST search.

IRESbase allows and encourages users to submit novel experimentally validated IRESs. In the submission, several basic IRES information (*e.g.*, IRES sequence, organism, host gene ID, host gene name, and host transcript ID) and validation assay information (*e.g.*, bicistronic backbone, positive and negative controls, second cistron expression, tested cell types, methods to analysis RNA structure, *in vitro* translation experiments, and PubMed ID) should be provided. In addition, users can also submit novel verified IRES by emailing a PageTab format file, in which all the essential information should be assembled. The submitted record will be included in the next release after manual check by our curators.

### Comparison with existing IRES databases

The current version of IRESbase contains the largest collection of experimentally validated IRESs in 11 eukaryotic organisms and 198 viruses. When compared with the existing IRES databases, IRESbase contains the largest number of eukaryotic IRESs and viral IRESs, including 1328 IRESs, while Rfam, IRESdb and IRESite only contains about 32 IRESs, 80 IRESs, and 135 IRESs in total, respectively (**Figure 5A**). In particular, IRESbase is the first database containing IRESs found in circRNAs and lncRNAs (Figure 5C). Moreover, IRESbase has minimal data redundancy. Unlike the IRESite database which collects all the truncated overlapping 5′ UTR sequences validated to contain IRES activity, IRESbase only stores the shortest functional sequences as the core IRES region. The median length of IRESs in IRESbase is 174 nucleotides, which is much shorter than that in IRESite (Figure 5B).

**Figure 5.**
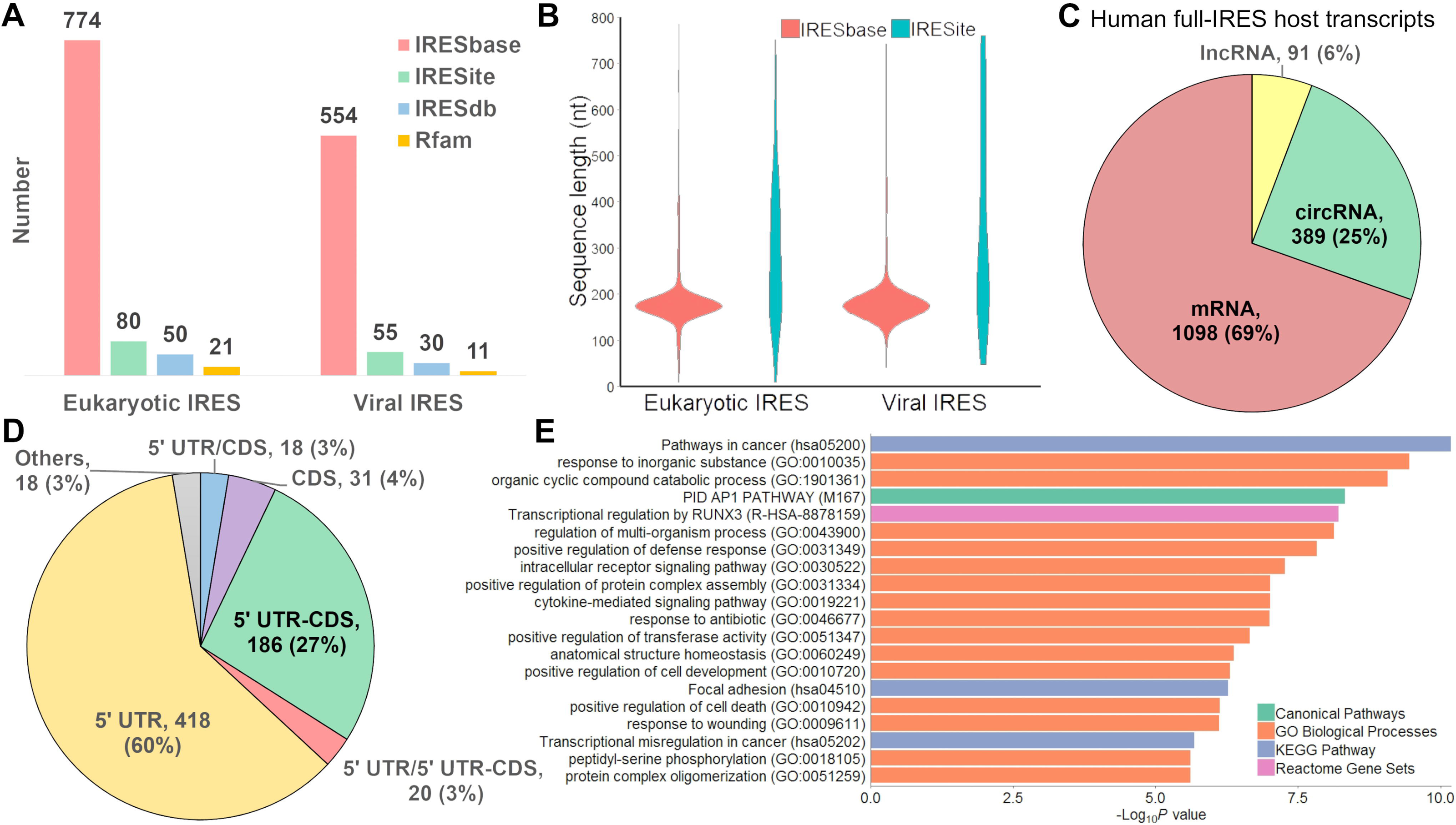
Statistic and analysis of human IRESs. **A.** Comparison with other IRES databases based on the number of IRESs curated. **B.** Comparison with IRESite based on the IRES length. **C.** The statistics of human full-IRES host transcripts (mRNA, lncRNA, and circRNA). **D.** The statistics of human IRES location in its host mRNA. **E.** The pathway and biological process enrichment analysis for the human IRES host genes (colored by enriched term).

In addition, IRESbase provides richer information and more useful functions than other IRES databases. Besides the experimentally verified 2D structure of 56 IRESs, IRESbase collected 12 IRES 3D structure and predicted 2D structure for all of the IRESs with Vienna RNA package [33]. It also provides more annotations, including genomic positions, sequence conservation, SNPs, nucleotide modifications, and targeting microRNAs for human IRESs. And apart from the IRES host transcripts collected from literature, IRESbase also provides many other potential IRES host transcripts (mRNA, lncRNA, and circRNA) predicted by sequence identity. Compared with existing databases, IRESbase provides more query fields (*e.g.*, genomic location, gene synonym) and search methods (fuzzy, advanced, and BLAST search). Additionally, the entire IRES dataset or those matching specific terms can be easily batch downloaded from IRESbase.

### Data statistics and analyses

In addition to 5′ UTRs, IRESs were also found in the CDS region. In the 691 human IRESs, 60% IRESs were found to be located only in the 5′ UTR region, 27% IRESs crossed the start codon, and 4% IRESs were only in CDS regions (Figure 5D). To analyse the functions of human IRESs, the gene set enrichment analysis was performed for the human IRES host genes using Metascape (www.metascape.org) [34]. As shown in Figure 5E, the most significant term is ‘hsa05200: Pathways in cancer.’ The enriched terms indicate that human IRESs may play important roles in cancer, cellular stress response, virus infection, and cell signalling.

As early as 1995, IRESs in mRNAs have been shown to be effective in initiating the protein synthesis of artificial circular RNAs [35], and in recent studies, IRESs found in circRNAs were also proved to be able to initiate the translation of linear transcripts (*e.g.*, bicistronic transcripts) [19–23]. All of the above-mentioned reasons indicate that the activity of IRES is not specific for the RNA type and IRES can play roles in any kinds of host RNAs. As only nine circRNAs and one lncRNAs have been respectively found to be translated via IRES so far, in order to find more potential coding circRNAs and lncRNAs, mRNA-derived IRESs were mapped to circRNA and lncRNA by using the pairwise sequence alignment. Complete 167 IRES sequences derived from mRNAs were found to be located within 371 circRNAs, and 45 complete IRES sequences derived from mRNAs were found in 83 lncRNAs. In particular, three complete IRES sequences derived from circRNAs were found to be located in eight mRNAs. Moreover, 1777 circRNAs and 77 lncRNAs were found to contain partial sequence (at least 30 nucleotides) of mRNA derived IRESs.

In order to investigate the potential coding circRNAs and lncRNAs, open reading frames (ORFs) were predicted for the circRNAs and lncRNAs containing a full or partial IRES sequence. Of the 371 circRNAs with an entire mRNA-derived IRES element, 261 were found to contain an ORF (> 300 nucleotides). Further, of the 1777 circRNAs with a partial mRNA-derived IRES element, 722 circRNAs were found to contain an ORF (> 300 nucleotides). The lncRNAs were all predicted to contain small ORFs (longer than 60 nucleotides and shorter than 300 nucleotides). And in 61 lncRNAs, the IRESs were found to be located between two small ORFs, which indicated that these lncRNAs may code multiple small peptides. In addition, the coding potential score was calculated for the circRNAs and lncRNAs, and the results suggested that 779 circRNAs and three lncRNAs were able to code proteins and small peptides, respectively.

## Discussion and perspectives

In recent years, as a functional RNA element which recruits ribosome to initiate translation in a cap-independent manner, IRES was found to mediate circRNA and lncRNA translation [18,24]. Although only a few peptides coded by the circRNA have been found, these peptides were found to play important roles in various biological processes [19–23]. In addition to the ORF, it was found that the coding ability of circRNA further depends on the IRES element. However, so far only a few IRESs are available in existing IRES databases, and the translation initiation mechanism via IRES is still elusive, especially for cellular IRESs. Therefore, there is an urgent need for a comprehensive curated dataset of all experimentally validated IRES elements, which could be a rich resource for functional studies and provides clues for circRNA and lncRNA protein coding ability.

In the near future, we will collect and update more IRES related information, including regulatory proteins, critical sites or regions for their activity, and experimentally verified biological function. We also plan to expand the database with experimentally validated sequences without IRES activity, which can not only help prevent repeated validations, but also facilitate the development of bioinformatics tools for the identification of IRES elements. Finally, as an open-access IRES database, we hope researchers can not only use its content but also help us update the database with information on new IRESs and provide us with a feedback.

## Materials and methods

### IRES dataset

To construct IRESbase with high quality information, the experimentally validated IRES sequences were manually searched from literature published before Oct. 14, 2019 using PubMed with either of the keywords “IRES,”, “IRESs”, “internal ribosome entry site”, “internal ribosome entry sites”, “internal ribosomal entry site”, “internal ribosomal entry sites”, “internal ribosome entry segment”, “internal ribosomal entry segment”, “internal ribosome entry sequence”, and “internal ribosomal entry sequence” in the titles or abstracts (Figure S2). We manually examined the abstracts of the retrieved literature, further examined the corresponding full texts and validated a novel IRES through experiments. For the literature concerning previously discovered IRESs, we further found the original publications through their citations. For published reviews and databases of IRESs, we checked all the citations of their collected IRESs, and located the original publications which discovered and validated these IRESs. The sequences of experimentally validated IRESs together with experiment information essential for the validation, including bicistronic backbone, positive and negative controls, second cistron expression, tested cell types, methods to analyse RNA structure, and *in vitro* translation experiments, were extracted. Instead of collecting all the truncated 5′ UTR sequences experimentally validated to contain IRES activity as IRESite did, only the shortest one was preserved in IRESbase to reduce data redundancy (Figure S3). In total, from 236 literature, 1328 experimentally validated IRES sequences, including 554 viral IRESs, 691 human IRESs, and 83 other eukaryotic (i.e., mouse, fly, yeast, *etc.*) IRESs, were collected.

### Collection of IRES sequences

As the available IRES sequence-related information are varied in different literature, we manually extracted the IRES sequence using different methods (Figure S2), which can be roughly classified into three categories: (a) extracted IRES sequence directly from the figure of its 2D structure, (b) extracted IRES sequence indirectly through the information of its host transcript and position, and (c) extracted IRES sequence by using the NCBI BLAST tool with the forward and reverse primers reported in the literature. Note that if more than one of the overlapping sequences was experimentally validated to contain IRES activity in literature, only the minimal functional one in them was selected (Figure S3). In addition, the IRES sequences derived from different literature were compared pair wise, and if the sequence identity was over 90%, only the shortest one was selected.

### Annotation of IRESs

We searched and collected the 2D structures of IRESs (or IRES host 5′ UTRs) from literature. The 3D structures of IRESs were manually searched and collected from the PDB, PDBj, and PDBe databases [36–38]. The sequences in the collected IRES 2D structures were manually extracted from the corresponding figures, and then the IRES-matched regions were calculated by sequence alignment. In addition, for all IRESs, the secondary structure and the minimal free energy (MFE) were also calculated using RNAfold of Vienna RNA package [33].

For human IRESs, genomic position, sequence conservation, single nucleotide polymorphisms (SNPs), RNA modification sites, potential host circular RNA, and microRNA-IRES interaction data were provided. As shown in Figure S4, the IRES genomic positions were obtained by mapping the IRES sequences to their host transcript sequences, and chromosome positions were extracted from NCBI transcript annotation files (GRCh37.p13 and GRCh38.p10) [39]. Based on the genomic positions, the phyloP scores (hg38 phyloP100way) were extracted from the UCSC database [28]. The dbSNP-NCBI database (150 release) was searched for the common SNPs and flagged SNPs within the IRES sequences [40]. The RMBase v2.0 database was searched for RNA modification sites (*e.g.*, N6-methyladenosines, N1-methyladenosines, pseudouridine modifications, and 2′-O-methylations) within human IRES regions [32]. The potential miRNA-IRES interactions were predicted by miRanda (version: aug2010) with a score threshold of 150 [41].

### Annotation of IRES host genes and transcripts

The host genes and transcripts of IRESs were collected from the literature, if available. For those IRES sequences without detailed host gene or transcript information in the literature, the nucleotide BLAST in NCBI was used to search for their host genes and transcripts with a sequence identity of at least 95% [42]. For viral IRESs, the closest gene with CDS downstream to the IRES element was regarded as its host gene. For human IRESs, besides mRNAs, we also aligned them to lncRNAs and circRNAs, and only those containing an IRES fragment of at least 30 nucleotides were considered as candidate IRES host transcripts, and 30 nucleotides were derived from the minimum length of known IRES elements.

For each IRES entry, the mRNA and lncRNA information were extracted from the NCBI nucleotide database, and the circRNA information was extracted from circBase, circRNADb, and circAtlas [25–27]. The ORF (< 300 nucleotides) and its corresponding peptide sequence were predicted for each lncRNA, and the ORF (> 300 nucleotides) and its corresponding peptide sequence were predicted for each circRNA. The coding potential score of ORFs was calculated by CPAT [43]. The sequence conservation of ORFs in lncRNA was assessed by conservation score, which was the mean phyloP score [28]. Gene level functional information (i.e., OMIM and Gene Ontology terms of biological process, molecular function, and cellular component) was extracted from the NCBI gene database, and the pathway information was retrieved from the KEGG PATHWAY database [30].

### Database construction

The IRESbase database was constructed by the assembly of all curated minimal IRES sequences and structure information, related transcript and gene information, and annotations. IRESbase was configured in a typical XAMPP (X-Windows + Apache + MySQL + PHP + Perl) integrated environment.

## Availability

IRESbase is freely accessible at http://reprod.njmu.edu.cn/cgi-bin/iresbase/index.php.

## Authors’ contributions

JZ designed the database structure, extracted IRES related information, constructed IRES datasets, and wrote the manuscript. YL searched and manually collected the IRES sequences from literature. CW and HTZ designed and constructed the website. HZ and BJ joined the study. XJG and XFS conceived the study, supervised the work, manuscript writing and editing. All authors read and approved the final manuscript.

## Acknowledgements

The study was supported by National Natural Science Foundation of China [No.61973155], the National Key R&D Program [2016YFA0503300], National Natural Science Foundation of China [No.61571223, No.81971439, and No.81771641], the Program for Distinguished Talents of Six Domains in Jiangsu Province [YY-019], the Fok Ying Tung Education Foundation [161037], the Fundamental Research Funds for the Central Universities [NP2018109], and Natural Science Foundation of the Jiangsu Higher Education Institutions of China Grant [18KJB310006], and the Scientific Research Foundation of Nanjing Medical University (2242019K3DN02).

## Supplementary material

**Figure S1 The homepage of IRESbase**

**Figure S2 The workflow of collecting experimentally verified IRES elements**

**Figure S3 The illustration for the selection of the minimal IRES**

**Figure S4 The illustration for the annotation of human IRESs**

