## Supplementary figures and images for "IRESbase: a Comprehensive Database of Experimentally Validated Internal Ribosome Entry Sites"

### Figure S1

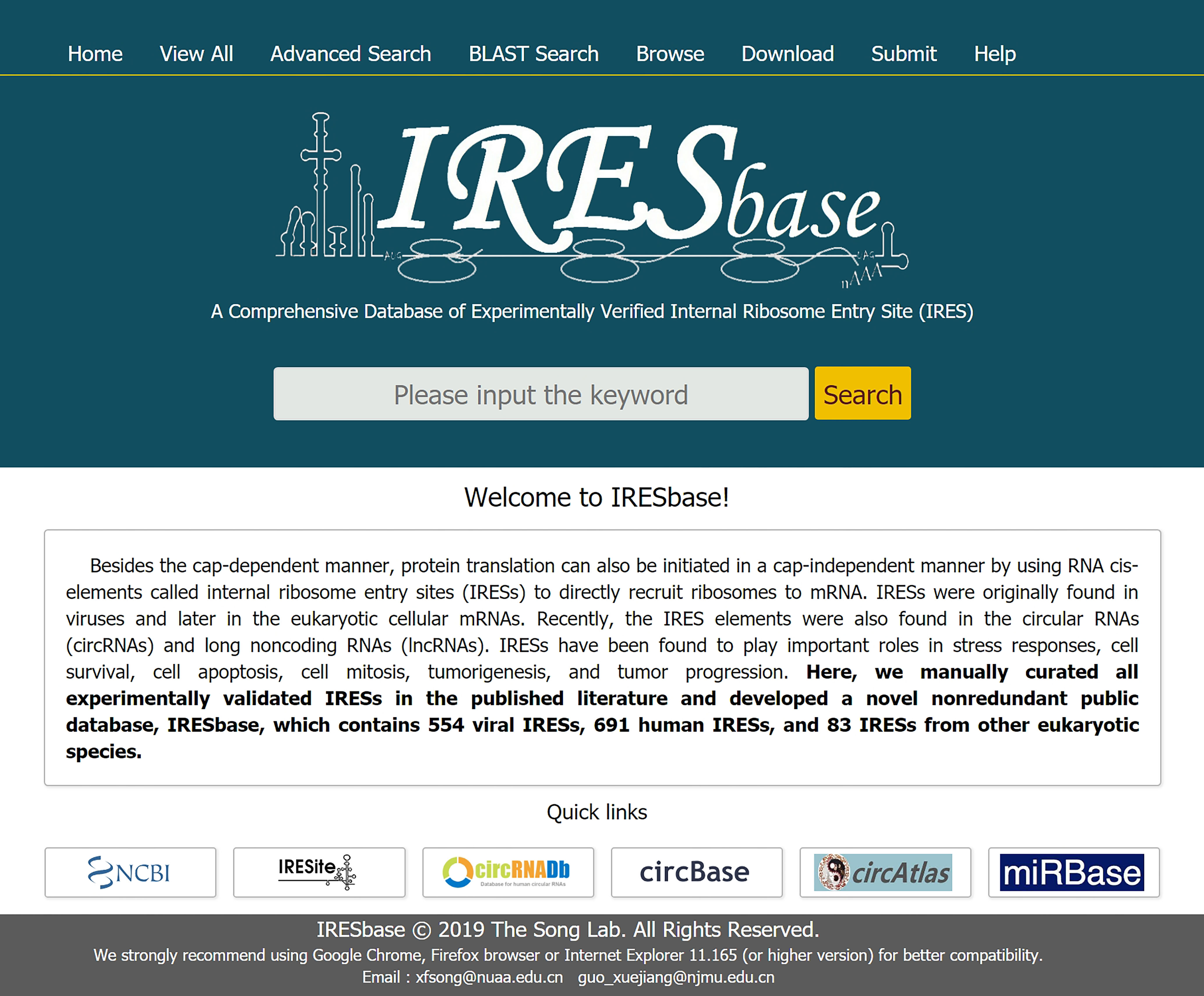

### Figure S2

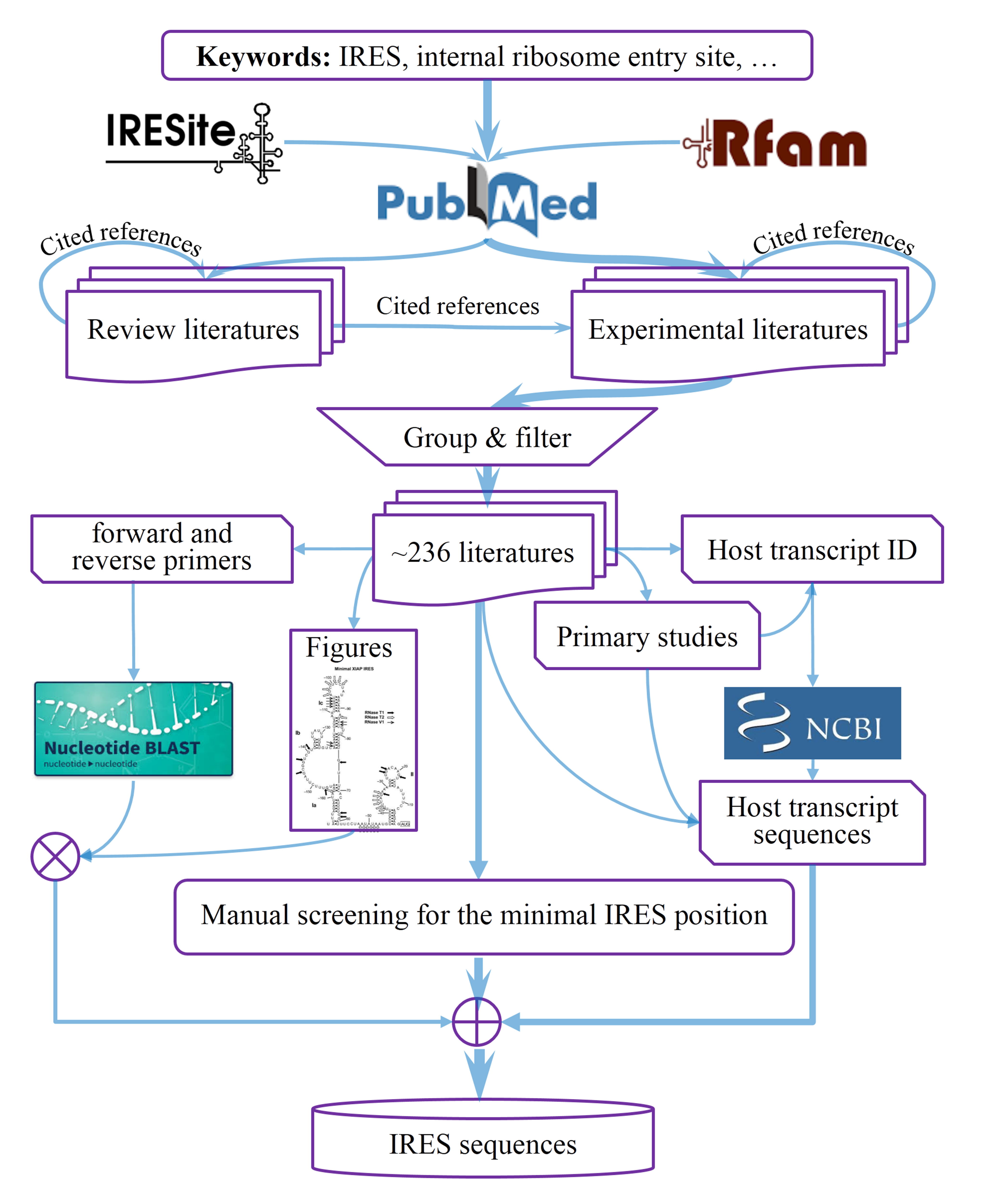

### Figure S3

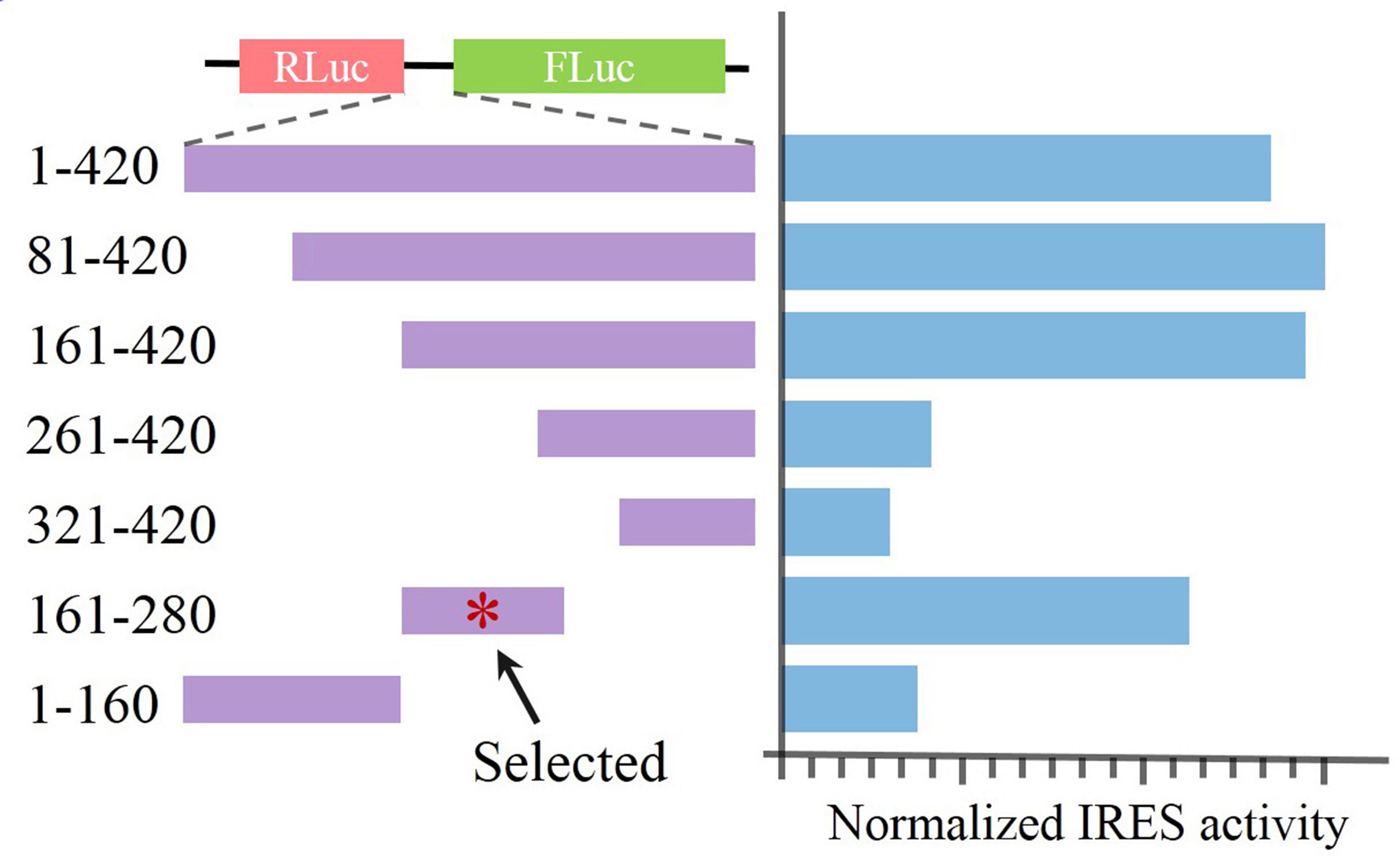

### Figure S4

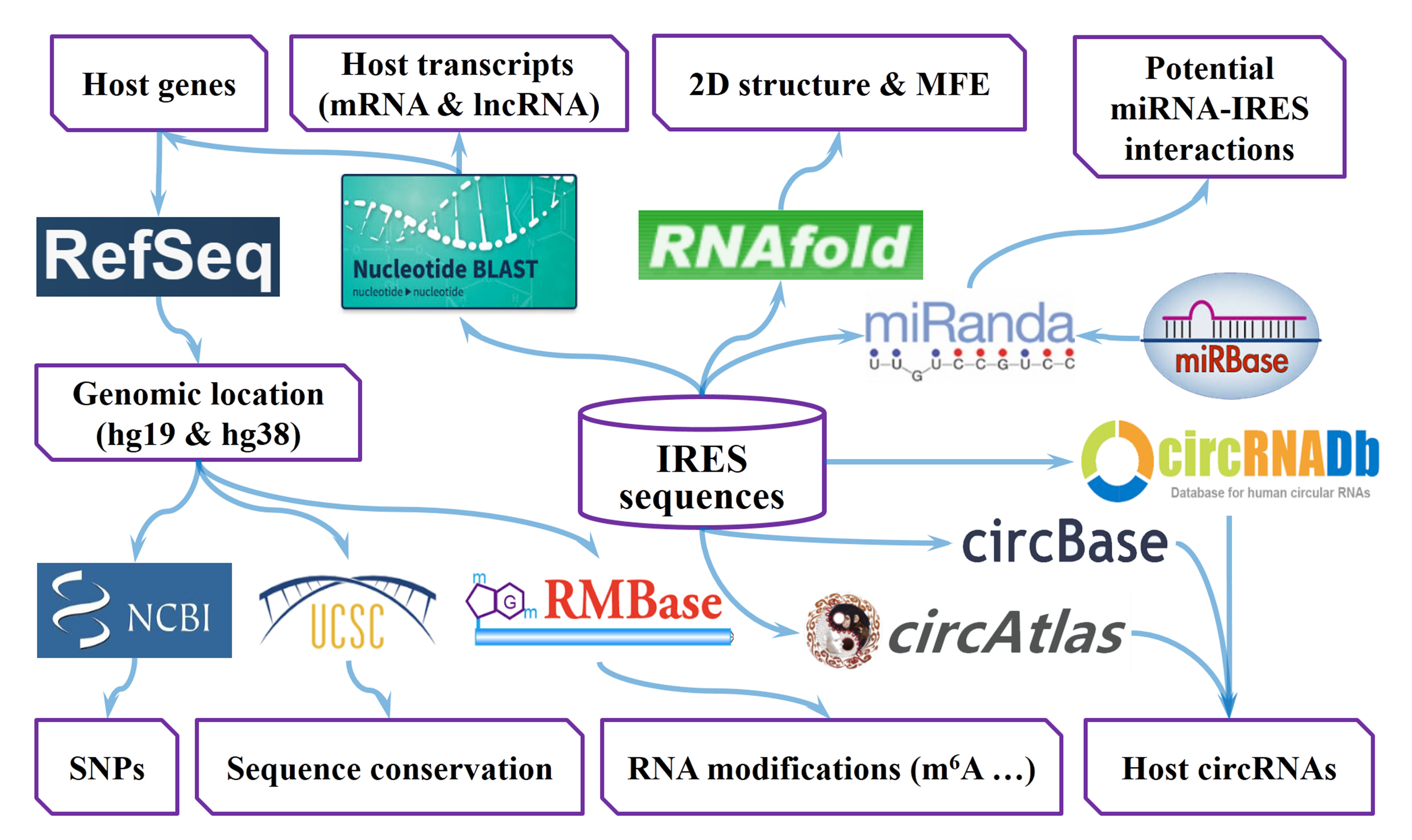
